# *APOE4* is associated with elevated blood lipids and lower levels of innate immune biomarkers in a tropical Amerindian subsistence population

**DOI:** 10.1101/2021.03.19.436070

**Authors:** Angela R Garcia, Caleb E Finch, Margaret Gatz, Thomas S Kraft, Daniel Cummings, Mia Charifson, Daniel Eid Rodriguez, Kenneth Buetow, Bret Beheim, Hooman Allayee, Gregory S Thomas, Jonathan Stieglitz, Michael Gurven, Hillard Kaplan, Benjamin C Trumble

## Abstract

In post-industrial settings, *APOE4* is associated with increased cardiovascular and neurological disease risk. However, the majority of human evolutionary history occurred in environments with higher pathogenic diversity and low cardiovascular risk. We hypothesize that in high-pathogen and energy-limited contexts, the *APOE4* allele confers benefits by reducing baseline innate inflammation when uninfected, while maintaining higher lipid levels that buffer costs of immune activation during infection. Among Tsimane forager-farmers of Bolivia (N=1266), *APOE4* is associated with 30% lower C-reactive protein, and higher total cholesterol and oxidized-LDL. Blood lipids were either not associated, or negatively associated with inflammatory biomarkers, except for associations of oxidized-LDL and inflammation which were limited to obese adults. Further, *APOE4* carriers maintain higher levels of total and LDL cholesterol at low BMIs. These results suggest the relationship between *APOE4* and lipids is likely beneficial for pathogen-driven immune responses, and unlikely to increase cardiovascular risk in an active subsistence population.

## 1. Introduction

The apolipoprotein *E4* (*APOE4*) allele is considered a major shared risk factor for both cardiovascular disease (CVD) (Hansson and Libby, 2006) and Alzheimer’s disease (AD) (Belloy et al., 2019; Smith et al., 2019), in part due to its role in lipid metabolism and related inflammation (Huebbe and Rimbach, 2017). *APOE4+* carriers consistently show higher levels of total cholesterol, low-density lipoprotein (LDL), and oxidized-LDL (Safieh et al., 2019, Yassine and Finch 2020). While some studies suggest *APOE4+* carriers have higher inflammatory responses (Gale et al., 2014; Olgiati et al., 2010), the *APOE4* allele is also associated with downregulation of aspects of innate immune function at baseline, including acute-phase proteins (Lumsden et al., 2020; Martiskainen et al., 2018; Vasunilashorn et al., 2011), and toll-like receptor (TLR) signaling molecules (Dose et al., 2018).

*APOE*, lipids, and immune function may interact differently in contemporary obesogenic post-industrial contexts compared to environments where infections are prevalent. For instance, a prospective U.S.-based cohort study (Framingham Heart Study) found that individuals with *APOE4* and low grade chronic obesity-related inflammation had higher risk of developing AD, with earlier onset than *APOE3/3* and *APOE4+* carriers without inflammation (Tao et al., 2018). By contrast, under high pathogen conditions, *APOE4* may protect against cognitive loss (Oriá et al., 2005; Trumble et al., 2017) and accelerate recovery from viral infection (Mueller et al., 2016). These findings suggest that interactive influences of *APOE4*, lipids, and immune function on disease risks may be environmentally mediated. However, this is difficult to test because most biomedical research is conducted in controlled laboratory settings using animal models, or in post-industrial populations with low pathogen burden and high obesity prevalence (Gurven and Lieberman, 2020). Here, we begin to fill this gap by evaluating both immune and lipid profiles of individuals with *APOE3/3* and *APOE4+* genotypes living in a high-pathogen, energy-limited environment.

### 1.1 *APOE*, cholesterols, and immune function

Despite its potentially deleterious health consequences, the ancestral *APOE4* allele is maintained in many human populations at nontrivial frequencies (as high as 40% in Central Africa) (Huebbe and Rimbach, 2017). A leading explanation for *APOE4* persistence is based on the theory of antagonistic pleiotropy (Williams, 1957), which posits that the *APOE4* allele may persist due to the fitness benefits of lipid buffering in early life relative to the *APOE3* mutation, outweighing any harmful health effects that manifest in a post-reproductive life stage (i.e “selection’s shadow”) (Smith et al., 2019). Consistent with the notion of early life fitness advantage, the *APOE4* variant is associated with lower infant mortality and higher fertility among rural Ghanaians experiencing high pathogen burden (Van Exel et al., 2017).

However, innate immune responses with fever and systemic inflammation are also energetically expensive (Muehlenbein et al., 2010), and cholesterol and other lipids are necessary for fueling these responses (Tall and Yvan-Charvet, 2014). In high-pathogen and energy-limited environments, where there may be persistent pathogen-driven immune activation, the ability to maintain peripheral cholesterol levels would presumably be a benefit throughout life (Finch et al., 2016; Gurven et al., 2016).

Despite *APOE4* being associated with neuroinflammation among those with AD, (Kloske and Wilcock, 2020), in ’healthy’ individuals the *APOE4* allele is associated with downregulated baseline innate immunity. Specifically, blood levels of C-reactive protein (CRP) in *APOE4+* carriers are lower in several post-industrial populations (Lumsden et al., 2020; Martiskainen et al., 2018), as well as the Tsimane, an Amerindian population in rural Bolivia (and focus of the present study) (Vasunilashorn et al., 2011). Moreover, among the Tsimane, *APOE4+* carriers had lower levels of blood eosinophils (Trumble et al., 2017; Vasunilashorn et al., 2011). Other studies have documented lower baseline levels of certain proinflammatory cytokines (e.g. IL-6, TNF-alpha) in *APOE4+* carriers (Olgiati et al., 2010), and downregulated expression of biomarkers mediating innate immune sensing (TLR-signaling molecules) (Dose et al., 2018).

Experimental studies also showed the associations of *APOE4* with heightened innate and complement inflammatory responses to lipopolysaccharide (LPS) stimulation in (Gale et al., 2014; Tzioras et al., 2019). Maintaining lower baseline levels of innate immunity may minimize the accumulative damage caused by low grade innate inflammation over the long term, while still enabling strong targeted immune responses to pathogens following exposure (Franceschi et al., 2000; Trumble and Finch, 2019).

In energy-limited, pathogenically diverse environments, *APOE4+* carriers may thus be better able to tolerate energetic costs imposed by infection by having higher concentrations of circulating lipids to fuel immune responses, while also minimizing damage from exposure to generalized systemic inflammation through downregulation of innate immune function. By contrast, in post-industrialized contexts, without the moderating influences of parasites on both cholesterol and immune functions, non-pathogenic stimuli (e.g. obesity) may be more likely to trigger systemic low-grade inflammatory pathways and, in the absence of a brake, lead to arterial and vascular damage and disease.

### 1.2 Hypothesis and Aims

We hypothesize that, in pathogenically diverse and energy-limited contexts, an *APOE4* variant is beneficial because it (a) minimizes damage caused by chronic innate inflammation, and (b) maintains higher circulating cholesterol levels, which buffer energetic costs of pathogen-driven innate immune activation. In post-industrial contexts, where there is a relative absence of diverse pathogens and thus reduced pathogen-mediated lipid regulation, coupled with an overabundance of calories, the effect of *APOE4* on circulating lipids may instead incur a cost. Lifestyle factors that promote obesity and excessive circulating lipids may lead to sterile endogenous inflammation (Trumble and Finch, 2019) that overshadows any beneficial effects of *APOE4* on immune function. Thus, in high-calorie, low-pathogen environments, the *APOE4* variant has greater potential to lead to hyperlipidemia and coincide with related inflammatory diseases (Figure 1).

**Figure 1.**
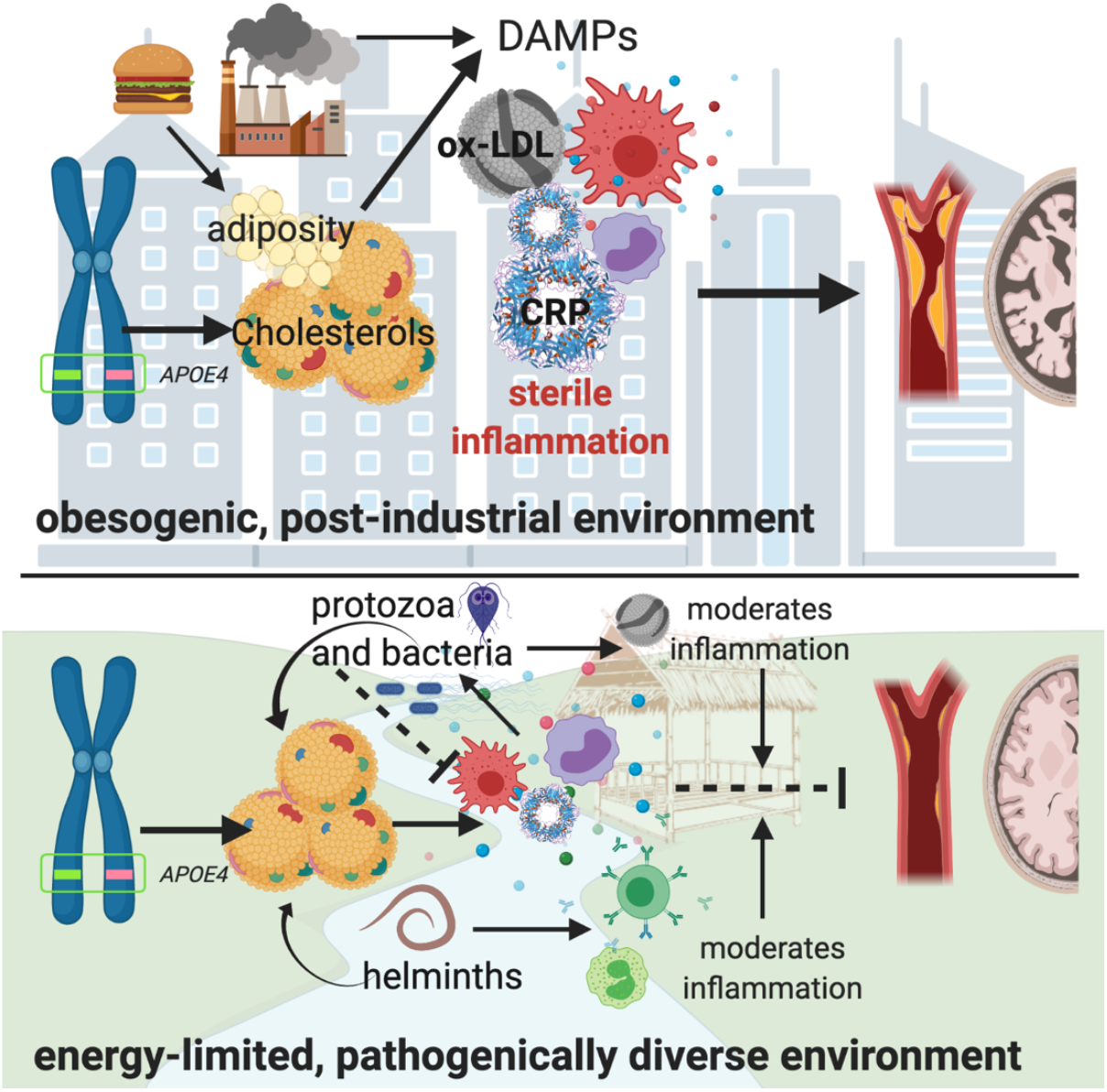
Hypothetical pathways through which the *APOE4* allele influences lipid processing, immune regulation and disease risk in post-industrialized and non-industrialized contexts. In both contexts, the *APOE4* allele leads to increased levels of circulating lipids; however, in post-industrialized contexts (a), lipid levels can reach dangerously high levels due to obesogenic diets, and an absence of moderation by parasites and pathogen-driven immune activation. Immune activation by non-pathogenic elements triggers damage-associated molecular pattern pathways, which generates a proinflammatory ’sterile’ immune response. Obesity and hyperlipidemia can simultaneously fuel sterile inflammation and promote oxidization of cholesterols, which, due to their lack of function, cause further tissue damage associated with cardiovascular and neurodegenerative disease risk. In energetically-limited, pathogenically-diverse contexts (b), the pathway between *APOE4* and disease risk is considerably more complex. Briefly, immune responses to parasites and microbes require cholesterol, and there are both direct and indirect effects of different species of parasites which further regulate cholesterol production and utilization. In addition, anti-inflammatory immune responses are generated by ox-LDL (e.g. in response to bacteria and protozoal infections), and helminthic parasites, which balance the immune system’s overall response. It is possible that in contexts where there is higher pathogen diversity, an *APOE4* phenotype may be beneficial because it minimizes the damage caused by upregulated innate immune functions, while also maintaining higher cholesterol levels which would buffer the cost of innate immune activation due to infection, whereas in high-calorie, low-pathogen environments, the utility of having an *APOE4* allele may be muted, and the costs more severe. Image created with BioRender.com

This study analyzes the immunophenotypes and lipid profiles of individuals with *APOE3/3* and *APOE4* genotypes living in a high-pathogen, energy-limited environment. We then evaluate the extent to which body mass index (BMI) moderates the association between lipids and inflammation. Finally, we test whether the *APOE4* allele has a moderating effect on the relationship between BMI and blood lipids to evaluate the role of *APOE4* in the maintenance of stable lipid levels under energetic restriction or pathogen stress.

This research focused on the Tsimane, an Amerindian population in the Bolivian tropics that faces high exposure to a diverse suite of pathogens, and endemic helminthic infections. Tsimane have high rates of infection across all ages, with 70% helminth prevalence and >50% of adults with co-infections from multiple species of parasite or protozoan (Blackwell et al., 2015; Garcia et al., 2020). In most villages, there is little or no access to running water or infrastructure for sanitation (Dinkel et al., 2020). The Tsimane are primarily reliant on foods acquired through slash-and-burn horticulture, fishing, hunting, gathering and small animal domestication, supplemented with market goods (e.g. salt, sugar, cooking oil) (Kraft et al., 2018).

Tsimane are rarely sedentary, instead engaging in sustained low and moderate physical activity over much of their life course (Gurven et al. 2013), and have minimal atherosclerosis (Kaplan et al., 2017). However, with greater globalization and improvements in technology, the Tsimane are experiencing ongoing lifestyle changes; there is variation in participation in the market economy, and related variation in diet (e.g. access and uptake of processed foods) and activity level.

## 2 Results

Data include *APOE* genotype and >6,500 measurements of BMI and seven immune markers in 1266 Tsimane adults. Multiple measurements per individual are used to capture individuals’ average levels, and minimize the potential of single or few outliers driving results (See Supplemental Table 1 for sample sizes per biomarker measurement). The sample is 50% female, and includes individuals from 80 villages. The median±SD age of the sample is 52±12 years (range: 24 -94). Median±SD BMI is 24±3 for both sexes; Tsimane adults are relatively short (women: 150.5±4.8cm; men: 161.7±5.3cm) with low body fat (median body fat percentage for men and women is 18% and 26%, respectively). Prevalence of obesity is relatively low (10%). In general, blood immune biomarkers vary significantly across adult ages (Supplemental Table 2; Figure 2). Table 1 provides a full description of lipid and immune levels of individuals by *APOE* genotype.

**Figure 2.**
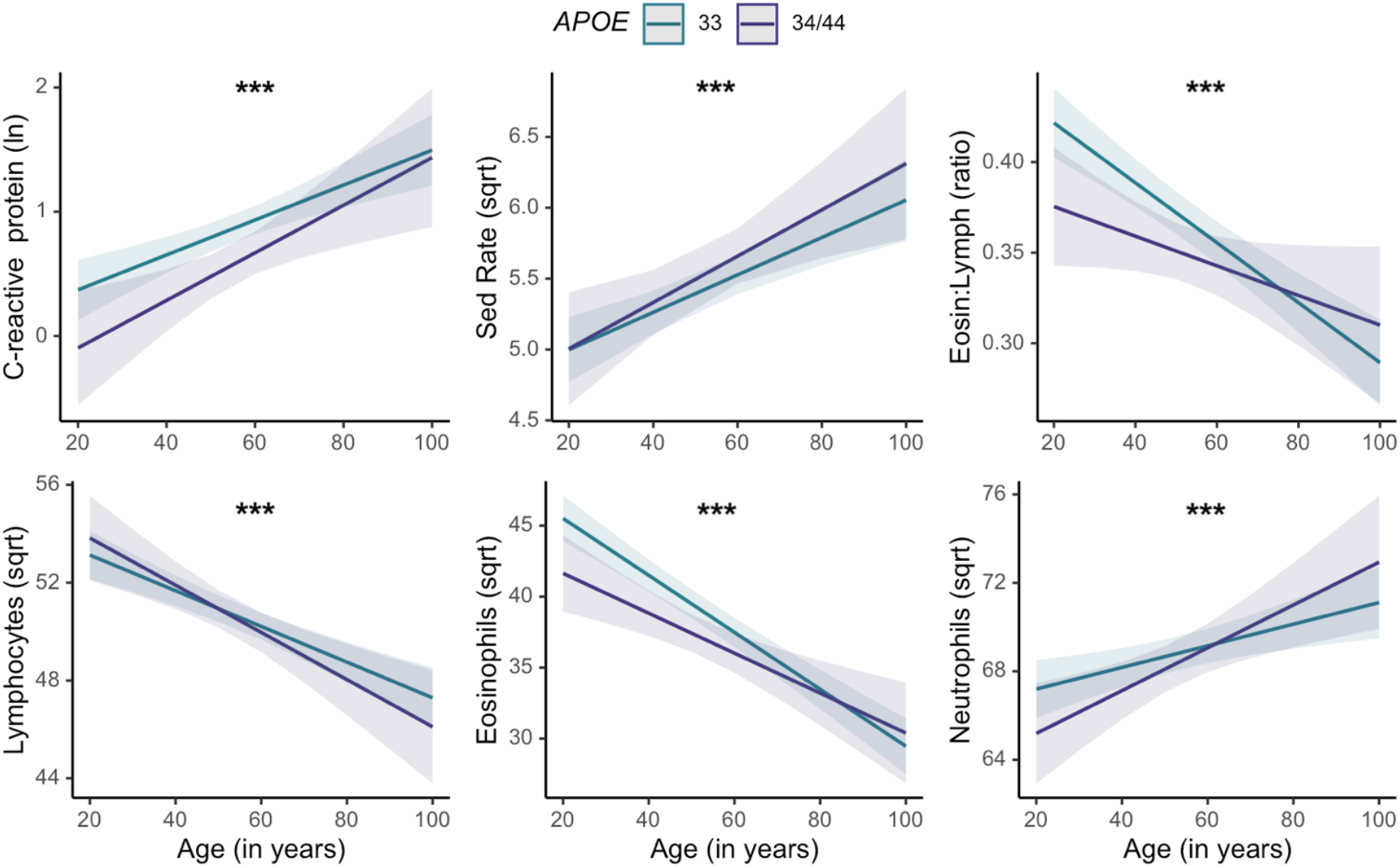
Plots showing estimated change in immune markers across age, split by *APOE* genotype. Slopes were taken from mixed effects linear regression models and represent estimates of the interactive effects of *APOE* genotype and age on each immune marker. Models adjust for sex, season, and current illness (Supplemental Table 2). Asterisks indicate significance level for overall differences in immune markers by age (all p<0.001). Erythrocyte sedimentation rate is abbreviated as ’Sed Rate’.

**Table 1.** Description of immune and lipid measures for homozygous *APOE3/3* and *APOE4+* carriers for whom age, sex, and BMI measures are available. Values are reported as mean (standard deviation). Linear mixed effects models fit by REML were used to test for differences between groups, controlling for age, sex, and seasonality, current infection, with random effects for community and individual. T-tests use Satterthwaite’s method. Due to skewness of biomarkers, statistical models use transformed and scaled data, for normalization. See methods for specific transformations for each marker.

| Variables | APOE 3/3 | APOE4+ | t-value | p-value |
| --- | --- | --- | --- | --- |
| N | 997 | 269 | --- | --- |
| % female | 51% | 48% | --- | --- |
| Age (in years) | 52 (12.1) | 52 (11.8) | 0.623 | 0.534 |
| BMI (kg/m<sup>2</sup>) | 23.8 (3.3) | 24.5 (3.6) | 2.215 | 0.027* |
| Hemoglobin (g/dL) | 13.2 (1.5) | 13.2 (1.4) | -0.768 | 0.442 |
| C-reactive protein (mg/dL) | 3.7 (3.3) | 2.9 (2.6) | -3.612 | <0.001*** |
| Leukocytes (1000/mm<sup>3</sup>) | 9.4 (2.6) | 8.9 (2.4) | -2.509 | 0.012* |
| Eosinophil (mm<sup>3</sup>) | 1618 (1061) | 1375 (910) | -3.789 | <0.001*** |
| Neutrophil (mm<sup>3</sup>) | 5045 (1803) | 4817 (1697) | -0.886 | 0.376 |
| Lymphocytes (mm<sup>3</sup>) | 2590 (832) | 2565 (852) | 0.153 | 0.878 |
| Eosin : Lymph | 0.36 (0.14) | 0.33 (0.14) | -3.149 | 0.002** |
| Sed Rate (mm/hr) | 29.5 (19.5) | 29.7 (19.2) | 1.262 | 0.207 |
| Triglycerides (mg/dL) | 107 (50.7) | 115 (61.0) | 1.097 | 0.273 |
| Total cholesterol (mg/dL) | 144 (32.3) | 148 (33.5) | 2.667 | 0.008** |
| HDL cholesterol (mg/dL) | 37.8 (8.8) | 38.0 (9.0) | 0.938 | 0.349 |
| LDL cholesterol (mg/dL) | 90.3 (32.2) | 92.9 (32.1) | 1.301 | 0.194 |
| Oxidized LDL (U/L) | 75.8 (24.0) | 78.6 (23.2) | 2.024 | 0.043* |

Notes: Statistical significance is denoted as: <sup>t</sup> $p<0.10$ ; \* $p<0.05$ ; \*\* $p<0.01$ ; \*\*\* $p<0.001$ .

In our sample, 21.2% have at least one *APOE4* allele; 245 individuals are heterozygous *3/4*, and 23 are homozygous *4/4*. The remaining 78.8% (n= 998) are homozygous for *APOE3/3*. Overall frequency of the *APOE4* allele is 12.7%. The *APOE2* allele was absent.

### 2.1 Characterization of lipid and immune profiles by APOE genotype

Relative to *APOE3/3* homozygotes, *APOE4* carriers have higher BMI (β=0.15 [CI: 0.02-0.28], p=0.03), total cholesterol (β=0.16 [CI: 0.04, 0.27], p=0.008), and oxidized-LDL (β=0.17 [CI: 0.01, 0.33], p=0.04), yet lower levels of innate immune blood biomarkers: CRP (β= -0.29 [CI: -0.45, -0.13], p<0.001), eosinophils, (β= -0.15 [CI: -0.23, -0.07], p<0.001), and a lower eosinophil to lymphocyte ratio (β= -0.02 [CI: -0.03, -0.01], p=0.002) (Table 1, Figures 3a and 3b). *APOE4* is also associated with lower total leukocytes (β= -0.09 [CI: -0.15, -0.02], p=0.01), but not with LDL, HDL, or triglycerides. Full models shown in Supplemental Tables 3 and 4.

**Figure 3.**
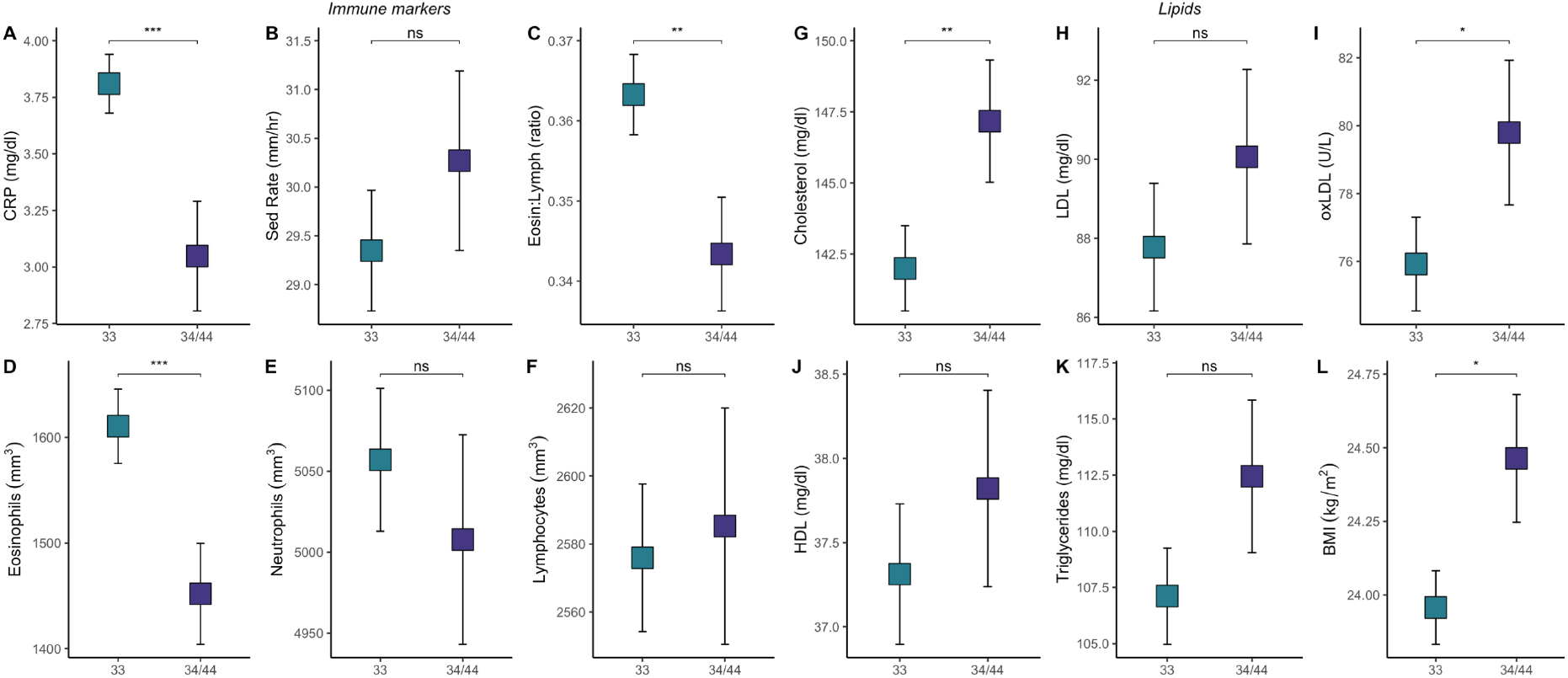
Barplots represent estimated means and standard deviations of immune and lipid markers for individuals that are homozygous *APOE* 33 versus those that have at least one copy of the *E4* allele. Estimates are standardized betas from mixed effects linear regression models that adjust for age, sex, season, and current illness (For full models with covariates see Supplementals T/able 3 and 4). Erythrocyte sedimentation rate is abbreviated as ’Sed Rate’.

### 2.2 Does BMI moderate the association between lipids and inflammation?

For Tsimane with higher (>24) BMI, total cholesterol and LDL did not associate with CRP; however, for individuals with median (20 ≤ BMI ≤ 24) or low (< 20) BMI, higher total cholesterol and LDL associate with lower CRP (Figure 4). When considered as a continuous variable, BMI significantly moderates these associations (total cholesterol: β=0.126 [CI: 0.07, 0.18], p<0.001; LDL: β=0.159 [CI: 0.09, 0.21], p<0.001). BMI also interacts with ox-LDL (β=0.141 [CI: 0.08, 0.19], p<0.001) in predicting CRP; however, this relationship is distinct from the other lipids tested. For Tsimane with high BMI, ox-LDL positively associates with CRP, while for those with low BMI the inverse is true (Figure 4). For erythrocyte sedimentation rate (ESR), total cholesterol (β=0.04 [CI: 0.01, 0.08], p=0.008) and LDL (β=0.03 [CI: 0.00, 0.06], p=0.09) positively associate with ESR only among Tsimane with higher BMIs. After adjusting for multiple testing, relationships between cholesterols and CRP all remain significant (all FDR adj. p <0.001), as does the relationship between total cholesterol and ESR (FDR adj. p=0.02).

**Figure 4.**
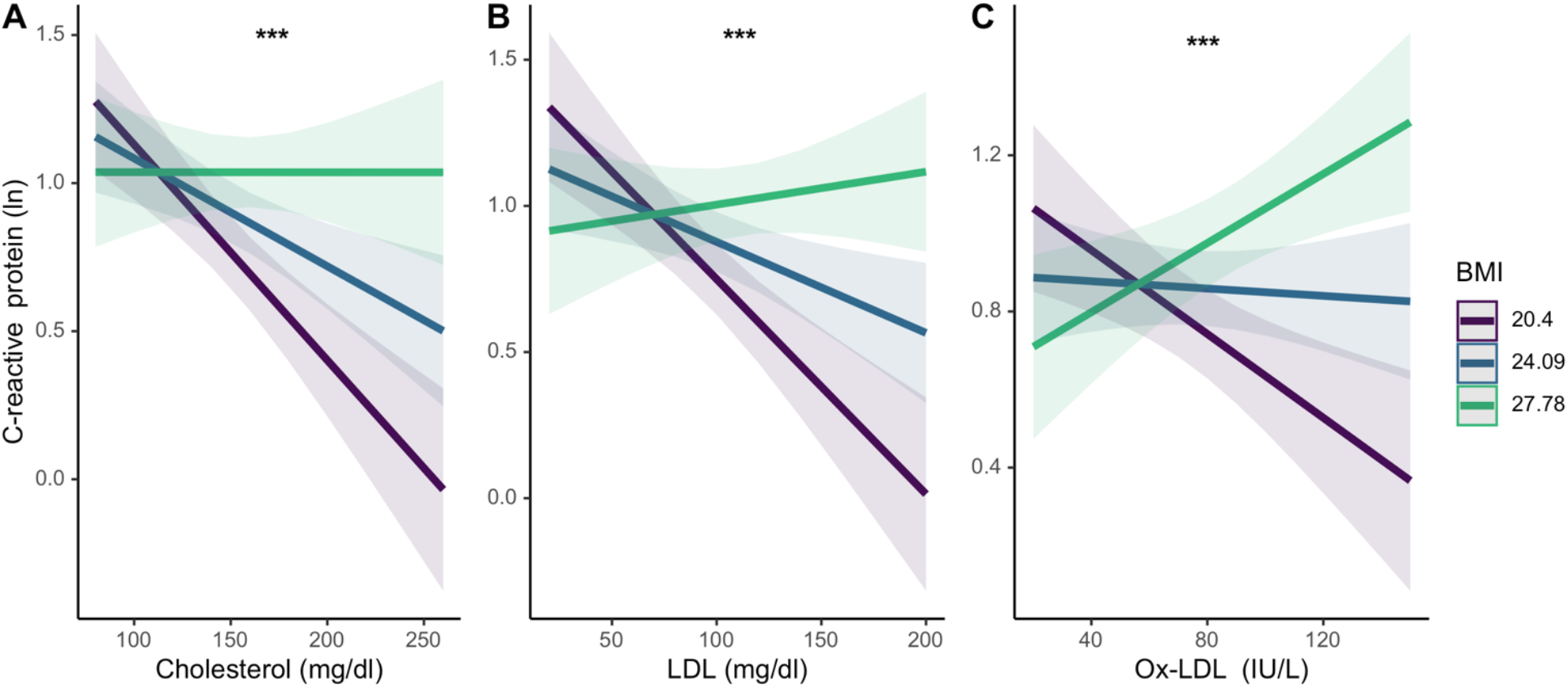
Differing influence of cholesterols on C-reactive protein, based on three levels of BMI (mean, ±1 standard deviation). Panels A-C show interaction effects between BMI and (A) total cholesterol; (B) LDL; and (C) oxidized LDL. For total and LDL cholesterol, among those with low (purple line) and mean (teal line) BMI, cholesterol is negatively associated with C-reactive protein. Oxidized LDL is only associated with higher CRP among individuals with high BMI (light green line). Results are reported as standardized betas from mixed effects linear regressions that adjust for age, sex, seasonality, with random effects for individual and community residence (Supplemental Tables 5-7). Variables are transformed and centered.

Concerning independent relationships, there are no direct relationships between LDL or ox-LDL and CRP, whereas higher total cholesterol is associated with lower CRP (β= - 0.14 [CI: -0.21, -0.06], p=0.002) (Supplemental Figure 1). In separate models, both BMI and total cholesterol are negatively associated with ESR (BMI:β= -0.05 [CI: -0.08, - 0.02], p=0.001; total cholesterol: β= -0.06 [CI: -0.09, -0.02], p=0.003). There are no independent or interactive relationships between BMI, cholesterols, and neutrophils. For full models see Supplemental Tables 5-7.

### 2.3 Does *APOE* genotype moderate the association between BMI and lipids?

Finally, we assess whether *APOE* genotypes differentially moderate associations between cholesterol and BMI for lean and obese individuals. To evaluate the effects of the *APOE4* allele on lipid levels across the range of BMI, we added an interaction term between BMI and *APOE* genotype to the mixed effects linear regression models assessing relationships between BMI and lipids (Table 2). These analyses show that *APOE4* carriers maintain similar levels of total cholesterol and LDL across BMIs (both: β = -0.04 [CI: - 0.07, -0.01], p=0.01), whereas *APOE3/3* homozygotes show higher cholesterol with BMI. Specifically, *APOE4* carriers maintain higher levels of total and LDL cholesterol at lower BMIs, but have lower levels of both at higher BMIs, relative to individuals that are homozygous *APOE3/3* (Figure 5). However, neither ox-LDL, HDL, nor triglycerides varied by *APOE* alleles across BMIs

**Table 2.** Models evaluating the moderating effects of APOE genotype on associations between BMI and cholesterols. Results are fixed effects estimates from mixed effects linear regressions, which include random effects for ID and community residence. In addition to age, sex, and season, a dummy variable was used as a proxy for current illness (leukocytes > 12 mm^3^). Results are reported as standardized betas; *CI* is the 95% confidence interval. All dependent variables were transformed and centered prior to analyses. *APOE* genotype is coded as a categorical variable, binned as individuals that are homozygous *E3* (E3) versus those that have at least one copy of the *E4* allele (E4).

| Predictors | Cholesterol |  |  | LDL |  |  | Ox-LDL |  |  | HDL |  |  | Triglycerides |  |  |
| --- | --- | --- | --- | --- | --- | --- | --- | --- | --- | --- | --- | --- | --- | --- | --- |
|  | B | 95% CI | p | B | 95% CI | p | B | 95% CI | p | B | 95% CI | p | B | 95% CI | p |
| *APOE* (*E4*) | 1.02 | 0.29 – 1.75 | **0.006** | 0.99 | 0.27 – 1.72 | **0.007** | 0.72 | -0.44 – 1.88 | 0.222 | 0.69 | -0.06 – 1.43 | 0.071 | 0.04 | -0.69 – 0.77 | 0.920 |
| BMI | 0.05 | 0.04 – 0.07 | **<0.001** | 0.06 | 0.04 – 0.07 | **<0.001** | 0.07 | 0.04 – 0.09 | **<0.001** | -0.03 | -0.04 – -0.01 | **<0.001** | 0.08 | 0.06 – 0.09 | **<0.001** |
| *E4* * BMI | -0.04 | -0.07 – -0.01 | **0.014** | -0.04 | -0.07 – -0.01 | **0.010** | -0.02 | -0.07 – 0.02 | 0.303 | -0.03 | -0.06 – 0.00 | 0.096 | -0.00 | -0.03 – 0.03 | 0.955 |
| Sex (male) | -0.13 | -0.22 – -0.04 | **0.005** | -0.17 | -0.25 – -0.08 | **<0.001** | -0.03 | -0.17 – 0.11 | 0.637 | 0.09 | -0.00 – 0.18 | 0.059 | -0.09 | -0.19 – 0.00 | 0.053 |
| Age (in yrs) | 0.01 | 0.01 – 0.01 | **<0.001** | 0.01 | 0.01 – 0.01 | **<0.001** | -0.00 | -0.01 – 0.00 | 0.200 | 0.01 | 0.00 – 0.01 | **0.002** | 0.01 | 0.00 – 0.01 | **0.006** |
| Season (wet) | -0.08 | -0.17 – 0.02 | 0.105 | -0.22 | -0.32 – -0.12 | **<0.001** | -0.31 | -0.50 – -0.11 | **0.002** | 0.10 | -0.00 – 0.20 | 0.054 | 0.13 | 0.04 – 0.21 | **0.003** |
| Currently ill | -0.03 | -0.20 – 0.13 | 0.689 | -0.06 | -0.23 – 0.11 | 0.507 | 0.07 | -0.22 – 0.37 | 0.633 | -0.14 | -0.32 – 0.03 | 0.105 | 0.02 | -0.13 – 0.17 | 0.797 |
| Intercept | -1.80 | -2.24 – -1.36 | **<0.001** | -1.78 | -2.21 – -1.34 | **<0.001** | -1.19 | -1.90 – -0.48 | **0.001** | 0.09 | -0.36 – 0.53 | 0.696 | -2.16 | -2.60 – -1.73 | **<0.001** |

**Figure 5.**
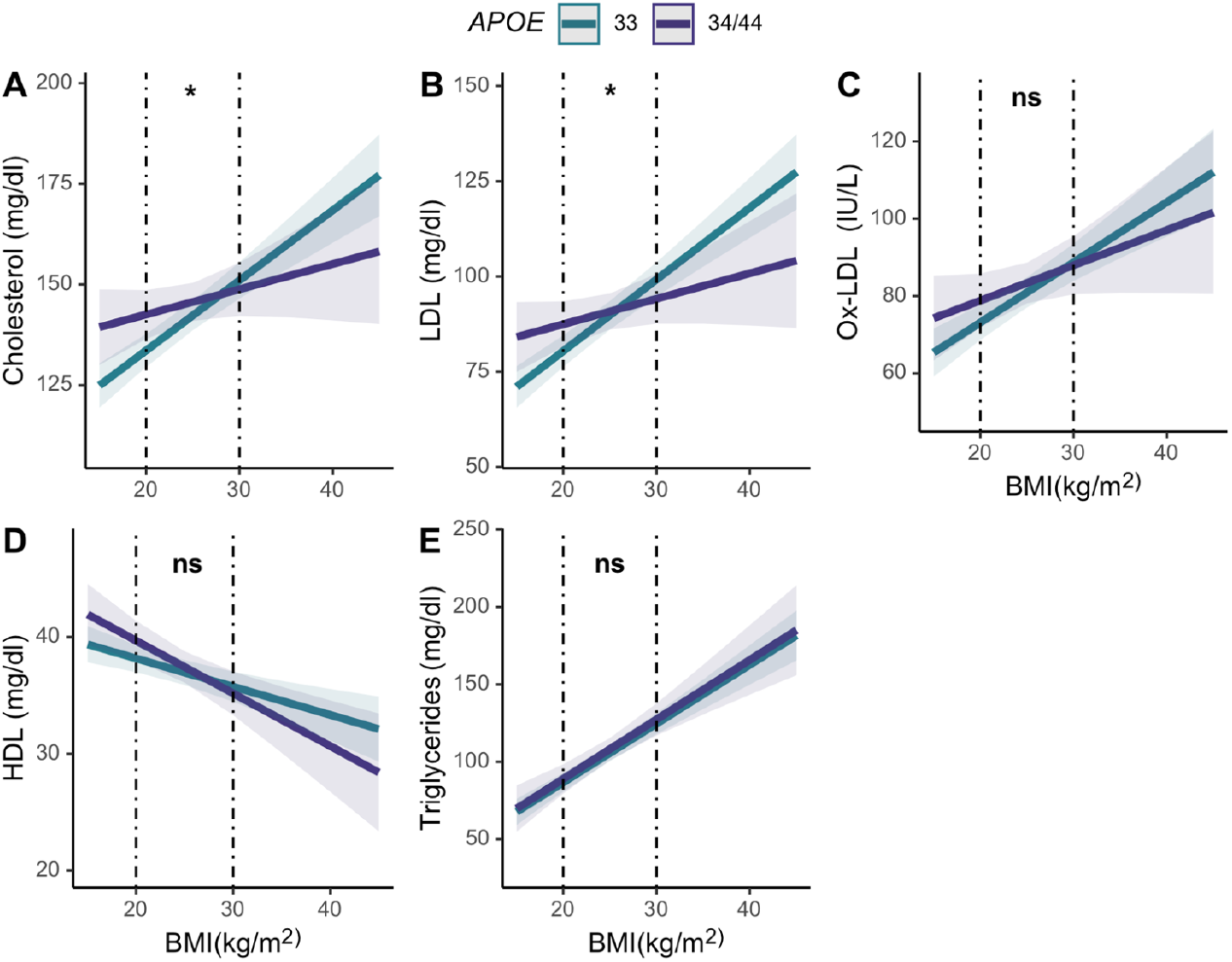
Plots showing moderating effects of *APOE* genotype on associations between BMI and cholesterols (from Table 2 in the manuscript). Dotted vertical lines represent cutoffs for low (<20) and high (>30) BMI. As predicted, for the primary cholesterols utilized during an immune response to pathogens-total cholesterol (A) and LDL (B)-individuals with an E4 allele maintain higher levels of those cholesterols at low BMIs, compared to E3 carriers. There were no significant differences across BMI for oxidized LDL, HDL, or triglycerides. Plots were derived from mixed effects linear regressions, and include age, sex, season, and current infection as covariates, as well as random effects for individual and community residence.

## 3 Discussion

We hypothesized that the effect of the *APOE4* polymorphism on disease risk may be environmentally-mediated, and that in an energy-limited, high-pathogen context, carrying an *APOE4* allele may be beneficial if it aligns with downregulated baseline innate immune function and higher circulating lipids. In support of this hypothesis, we find that Tsimane with an *APOE4* allele have higher levels of lipids, yet lower levels of C-reactive protein and eosinophils, compared to individuals that have a homozygous *APOE3* genotype. Overall, *APOE4* carriers also have an adaptive-biased immune profile (as measured by a lower ratio of eosinophils to lymphocytes). These associations remain when controlling for factors known to contribute to differences in inflammation including age, sex, seasonality, and current infection. The most striking example of immunological differences is for CRP, an acute phase protein that functions as part of the Complement system and is a marker of generalized inflammation. Though CRP is significantly higher in the Tsimane relative to U.S. and European populations (Blackwell et al., 2016; Gurven et al., 2008), we find that CRP levels are 30% lower among *APOE4* carriers than individuals that are homozygous *APOE3/3*. This replicates a similar finding of lower CRP among Tsimane *APOE4* carriers from samples collected over a decade prior to the current study (Vasunilashorn et al., 2011). A study of ’healthy’ (nondiabetic) Finnish men also found that *APOE4* carriers had higher LDL cholesterol coupled with lower CRP (Martiskainen et al., 2018).

Further, in support of our hypothesis that lifestyle and ecological (e.g. pathogen exposure) factors affect how lipids influence inflammation, we find that BMI significantly moderates relationships between lipids and markers of innate inflammation. Our results are consistent with expectations for a high infection context: Tsimane with high BMI (∼1 SD above the mean) show no relationship between total cholesterol or LDL with CRP and erythrocyte sedimentation rate (ESR), but those with mean or low BMI have *higher* lipid levels that are associated with *lower* CRP and ESR. In contemporary post-industrialized contexts (Fig. 1a), these findings may seem counterintuitive. But in a high pathogen context, higher concentrations of circulating lipids may allow individuals, especially those that are energy-compromised, to better tolerate infection (Gurven et al., 2016). Thus, the negative association between cholesterol and inflammatory markers could indicate lower infectious burden. For the Tsimane, higher cholesterol levels are likely an indicator of robust health (e.g. not fighting an infection) rather than chronic disease. Independently, lipids are either neutral or negatively associated with immune markers in this sample. This is consistent with previous research among the Tsimane that found blood lipids varied inversely with IgE, eosinophils, and other markers of infection (Vasunilashorn et al., 2010).

Notably, the moderation effect of BMI indicates that the relationship between ox-LDL and CRP are inverted at the low and high ends of the BMI range. For individuals with low BMI, higher ox-LDL levels are associated with lower CRP, while for those with high BMI, higher ox-LDL is associated with higher CRP (Figure 4c). One possible explanation for the opposing relationships we find between ox-LDL and CRP is that there may be differences in the underlying causes of oxidization in these groups. Though ox-LDL is often considered a consequence of obesogenic or hyperlipidemic oxidization of LDL (Neuparth et al., 2013), ox-LDL is also produced in response to pathogen-mediated immune activation (Han, 2009), and its relationship with downstream immune functions largely depends on the cause of oxidation. For example, while nonpathogenic oxidization of LDL promotes inflammation (Neuparth et al., 2013; Tall and Yvan-Charvet, 2014), during protozoal or bacterial infections, oxidized LDL contributes to lower inflammatory responses (Han, 2009). Thus, those with high BMI may represent the transition to sterile inflammation as lifestyles change with greater participation in a market economy (Trumble and Finch 2019). Indeed, other studies have documented what may be the start of a nutritional transition among the Tsimane (Kraft et al., 2018; Masterson et al., 2017; Rosinger et al., 2013).

Our finding that innate immune biomarkers are lower among *APOE4* carriers is in line with prior reports (Lumsden et al., 2020; Martiskainen et al., 2018; Trumble et al., 2017; Vasunilashorn et al., 2011), however the causes are uncertain. One proximate explanation involves the mevalonate pathway, which plays a key role in multiple cellular processes, including modulating sterol and cholesterol biosynthesis (Buhaescu and Izzedine, 2007). The current study design did not allow analysis of this pathway.

Finally, though numerous other studies have established links between the *APOE4* polymorphism and neighboring genes (the *APOE* gene cluster) (Kulminski et al., 2019), and increased circulating lipids (Yassine and Finch 2020, Posse de Chaves and Narayanaswami, 2008; Safieh et al., 2019; Saito et al., 2004), to our knowledge, this study is the first to assess whether *APOE* moderates lipid levels under energetic constraints. Specifically, our results show that the *APOE4*+ genotype is associated with higher relative lipids in a low-energy, high-pathogen system. In one study conducted in an obesogenic environment (mean BMI 28 kg/m^2^), BMI was independently associated with higher total cholesterol, but this relationship did not differ by *APOE* genotype (Petkeviciene et al., 2012). While it is difficult to pinpoint the cause for discrepant findings across studies, one possibility is that the Lithuanian study did not capture moderation at low BMI (the mean BMI was 28 for both *E4 APOE4*+ and *APOE3/3* genotypes). Another possibility is that the low-pathogen context of urban Lithuania led to less moderation of cholesterols at high BMIs. As mentioned in section 1, one proposal for the persistence of the ancestral *E4* allele despite its deleterious health effects at later ages relies on the fitness benefits of lipid buffering in early life relative to the more recent *E3* mutation (Van Exel et al., 2017; Yassine and Finch, 2020). The ability to maintain adequate lipid reserves under energetic and pathogenic pressures would also provide additional benefits over the life course (Finch and Kulminski, 2020).

The commonly reported associations between *APOE4* and disease risk in post-industrialized societies may differ from subsistence populations due to environmental mismatch (Trotter et al., 2011). In an environment where calories are limited, and one’s innate immune system is already primed by multiple pathogens, the benefit of having an *APOE4* allele - which facilitates upregulated lipids (sufficient for mounting immune responses) and downregulates innate immune function - may be amplified. Such a phenotype would be beneficial particularly if downregulated baseline immune function did not compromise responses to immunological threats.

## 4 Limitations

Though our findings draw from a large sample size and are robust to various controls and model specifications, there are several limitations. First, our findings are correlative and limit causal inference. Because these findings may be important for furthering evolutionary (i.e. why the *APOE4* allele is maintained) and clinical (i.e. the role of *APOE* in disease pathogenesis) understanding, they require replication, and warrant experimental testing. The central thesis presented here – that persistent exposure to pathogens and obesogenic diets moderate the relationship between blood lipids and inflammation – is amenable to experimental manipulation under lab conditions. Specifically, a mammalian model system could be split into two treatments: those raised under sterile conditions versus regimented exposure to non-lethal pathogens. These treatments may then be crossed with dietary or physical activity conditions that produce differential levels of adiposity. Our hypothesis predicts that both decreased adiposity and increased life course pathogen exposure will reduce or even eliminate positive associations between blood lipids and chronic inflammation. Importantly, inflammatory biomarkers can be measured at more frequent intervals in lab conditions to assess long-term differences in the function of both pro- and anti-inflammatory pathways between experimental treatments.

Secondly, there is some evidence that the *APOE4* allele is positively associated with HDL cholesterol levels (Hopkins et al., 2002), and that higher HDL levels reduce risk of severe infection (measured by infectious hospitalizations) (Trinder et al., 2020). Inversely, acute infections and systemic inflammation (e.g. acute phase reaction) are associated with a decrease in HDL and HDL remodeling that results in lower cholesterol efflux capacity and higher peripheral levels of LDL cholesterol (Ronsein and Vaisar, 2017; Zimetti et al., 2017). While we did not find differences in HDL by *APOE* status among the Tsimane, we cannot completely discount the possibility that HDL remodeling plays a role in the higher lipid levels, in addition to *APOE* allelic variation. Further, given the relative lack of, and difficulty in accessing, medical care facilities, it is difficult to assess degrees of infection severity, and thus it is also possible that *APOE4* may mitigate infectious disease burden. However, the current data cannot provide evidence for either of these potentials, and further research is needed to disentangle the roles of *APOE* and lipids in infection.

Third, because patterns of immune response vary depending on pathogen type and species, the use of a high white blood cell count cutoff as a proxy for current infection overly simplifies immune variation due to different types of infection. However, we also adjust for seasonality and cluster by community residence in our models, which should capture additional variation in exposures.

Finally, we were not able to fully adjust for the time of day that samples were collected for the biomarkers used in this paper. Given that peripheral levels of most immune biomarkers vary diurnally to some extent, it is possible that not adjusting for exact time of day may have introduced some noise into analyses. However, CRP (which the main findings centered around) does not appear to follow a circadian rhythm in healthy individuals (Meier-Ewert et al., 2001). Further, the largest differences in levels (peak to trough amplitudes) tend to coincide with sleep and wake cycles (Labrecque and Cermakian, 2015). Because blood draws routinely occur in the morning, samples are constrained to a narrow window, and thus we are not comparing values across the full range of diurnal variation.

## 5 Conclusion

In post-industrial settings, *APOE4* is generally considered a purely deleterious allele, increasing inflammation and lipids, and escalating cardiovascular and neurological disease risk. Yet in a high pathogen environment with minimal obesity, we find that *APOE4* is associated with *lower* levels of innate inflammation. While *APOE4* carriers do have higher lipid levels, these are likely beneficial for immune response and child survival, and unlikely to increase CVD risk in a population without other cardiometabolic risk factors.

## 6 Materials and Methods

### Population and Sampling Design

Data come from the Tsimane Health and Life History Project (THLHP), a longitudinal study of health and behavior that has run continuously since 2002. The THLHP integrates methods from anthropology, epidemiology and biomedicine to better understand aging and the role of infection on chronic disease risk (Gurven et al., 2017).

The Tsimane are an Amerindian population that live in the tropical lowlands of Bolivia. As of 2015, the THLHP census estimated a total population size of about 16,000 individuals living across 90+ villages (Gurven et al., 2017). The Tsimane live a majority subsistence lifestyle, practicing slash-and-burn horticulture to cultivate crops (plantains, manioc, corn, rice) that comprise the majority of calories in their diet, and fishing, hunting, and collecting nuts, berries, and seasonal fruits (Kraft et al., 2018). Processed foods (e.g. sugar, salt, oil, flour) are becoming more common, though acquiring these items remains somewhat limited. Electricity and sanitary infrastructure are also severely limited, with most villages lacking clean water and access to items such as antiseptics and other antibacterial cleansers. From an early age, Tsimane are exposed to a diverse array of pathogens, and parasitic and other infections are common (Blackwell et al., 2016; Garcia et al., 2020; Vasunilashorn et al., 2010). Compared to U.S. and European reference populations, the Tsimane have also been found to have upregulated immunity across the life course (Blackwell et al. 2016). Despite increasing access to markets and towns, infections remain the largest source of morbidity (Gurven et al., 2020).

### Sampling Design

Biomarker data were collected by the THLHP (see Gurven et al., 2017; Kraft et al., 2020 for details). A Bolivian and Tsimane mobile medical team travel annually or biannually among study communities conducting clinical health assessments and collecting biochemical and anthropometric information from community members that want to participate. This sample includes all data from individuals for whom we have *APOE* genotyping and at least one measurement of BMI, sex, and age - which is the base criteria for this study. Sample size varies by biomarker and over time for several reasons: sampling strategy varies by data type, absent or sick team personnel needed to collect data, the number of study villages and thus enrolled participants has increased over time, and the data types collected have changed over time (see Kraft et al., 2020). Specific sample sizes per covariate are reported in Supplemental Table 1, and full tables report sample size for each model.

### Ethics

This research has been approved by institutional review boards at the University of New Mexico (#07-157) and University of California Santa Barbara (#3-20-0740), as well as the Tsimane government (Tsimane Gran Consejo) and village leaders. Study participants give consent for each part of the research and data collection prior to participating, during every visit by the THLHP.

### APOE genotyping

Whole blood samples were stored in cryovials (Nalgene, USA) and frozen in liquid nitrogen before transfer on dry ice to the University of California-Santa Barbara, where they were stored at -80°C until genotyping. Single nucleotide polymorphism (SNP) genotyping was used to identify *APOE* allelic variants in blood samples. Samples were shipped on dry ice to University of Southern California (2010 and 2013) and University of Texas-Houston (2016), where DNA was extracted, quantified, and haplotype coded for *APO-E2, E3*, and *E4* alleles using the TaqMan Allelic Discrimination system (Thermo-Fisher Scientific, Carlsbad, CA, USA). Determination of the *APOE2/E3/E4* alleles in the Tsimane derived from 2 SNPs of 20-30bp oligonucleotides surrounding the polymorphic site (Cys112Arg/rs429358 and Cys158Arg/rs7412) (Trumble et al., 2017; Vasunilashorn et al., 2011).

### Measurement of blood lipids and immune function

Biomarkers were either assayed in the field at the time of collection, or in the Human Biodemography laboratory at UC Santa Barbara in 2016.

Blood was collected by venipuncture in a heparin-coated vacutainer. Immediately following the blood draw, total leukocyte counts and hemoglobin were determined with a QBC Autoread Plus dry hematology system (QBC Diagnostics), with a QBC calibration check performed daily to verify QBC performance. Relative fractions of neutrophils, eosinophils, and lymphocytes were determined manually by microscopy with a hemocytometer by a certified Bolivian biochemist. ESR was calculated following the Westergren method (Westergren, 1957).

Serum was separated and frozen in liquid nitrogen before transfer to the University of California-Santa Barbara where a commercial immunoassay was used to measure oxidized LDL (Mercodia, Winston Salem, NC). Serum high sensitivity C-Reactive Protein (hs-CRP) was assessed via immunoassay (Brindle et al., 2010), and was cross-validated by the University of Washington laboratory, using the protocols utilized for the National Health and Nutrition Evaluation Survey (NHANES). Total and LDL cholesterol levels from serum samples were measured (Stat Fax 1908, Awareness Technology, Palm City, FL) in the THLHP laboratory in San Borja, Beni, Bolivia.

### Age estimation and anthropometrics

Birth years were assigned based on a combination of methods including using known ages from written records, relative age lists, dated events, photo comparisons of people with known ages, and cross-validation of information from independent interviews of kin (Gurven et al., 2007). Each method provides an independent estimate of age, and when estimates yielded a date of birth within a three-year range, the average was used. Individuals for whom reliable ages could not be ascertained are not included in analyses.

Weight and height were measured in the field by a member of the THLHP medical team, using a basic digital scale (Tanita, Arlington Heights, IL) and stadiometer to the nearest 0.1 cm. BMI was calculated as weight (kg) / height^2^ (m^2^).

### Statistical Analysis

Mixed effects linear regressions with restricted maximum likelihood estimation are used for all analyses. Models adjust for age, sex, seasonality, and current infection (leukocyte count > 12,000 cells per microliter of blood) (McKenzie and Williams, 2010), with random intercept effects for individual ID and community. *APOE* genotype is defined categorically, binning individuals as homozygous *APOE3* (*E3/3*) or as *APOE4+* carriers (if they had at least one copy of the *APOE4* allele). Heterozygous and *APOE4* homozygotes were binned together due to the small number of homozygotes. To model moderation effects (sections 2.2 and 2.3) interaction terms are included between the main predictor and moderator.

Immune and lipid measures required transformation to normalize their skewed distributions. Variables were transformed as follows: CRP, BMI, and triglycerides were natural log-transformed; total leukocytes and subsets (lymphocytes neutrophils, eosinophils), ESR, and remaining cholesterols (total cholesterol, LDL, HDL, ox-LDL) were square-root transformed. To compare across models, all dependent variables were then z-scored for analyses, and thus all betas are standardized estimates.

## 6 Acknowledgements

We thank Tsimane participants and their communities, and the THLHP field team (including administrators, logistical support, physicians, biochemists and anthropologists), whose support, expertise, and commitment made this work possible.

