## Supplemental Materials for "*APOE4* is associated with elevated blood lipids and lower levels of innate immune biomarkers in a tropical Amerindian subsistence population"

| **Variables** | **APOE 33** | **APOE 34/44** |
| --- | --- | --- |
| Total # of individuals (*N*) | 1002 | 271 |
| Age | 10183 (1000) | 2975 (269) |
| BMI | 5112 (998) | 1457 (268) |
| Hemoglobin | 5189 (998) | 1428 (268) |
| C-reactive protein | 925 (673) | 252 (175) |
| Leukocytes | 5171 (996) | 1422 (268) |
| Eosinophil | 4859 (993) | 1333 (268) |
| Neutrophil | 4871 (993) | 1337 (268) |
| Lymphocytes | 4862 (993) | 1335 (268) |
| Lymph : Eosin | 4859 (993) | 1333 (268) |
| Sed Rate | 4746 (986) | 1308 (268) |
| Triglycerides | 2330 (952) | 627 (257) |
| Total cholesterol | 2288 (949) | 616 (256) |
| HDL cholesterol | 2199 (950) | 585 (254) |
| LDL cholesterol | 2131 (942) | 567(254) |
| Oxidized LDL | 926 (673) | 252 (175) |

Supplemental Table 1. Longitudinal data used in analyses. split by APOE genotype. Table reports total # of measurements (at least one measurement per unique individual) for each marker.

|  | **CRP** | | **Sed Rate** | | **Neutrophils** | | **Eosinophils** | | **Eosin:Lymph** | |
| --- | --- | --- | --- | --- | --- | --- | --- | --- | --- | --- |
| *Predictors* | *β ( 95%CI )* | *p* | *β ( 95%CI )* | *p* | *β ( 95%CI )* | *p* | *β ( 95%CI )* | *p* | *β ( 95%CI )* | *p* |
| (Intercept) | -0.80 (-1.15 – -0.45) | **<0.001** | -0.29 (-0.45 – -0.12) | **0.001** | -0.33 (-0.47 – -0.19) | **<0.001** | 0.88 (0.72 – 1.04) | **<0.001** | 0.44 (0.41 – 0.46) | **<0.001** |
| *APOE4* | -0.29 (-0.45 – -0.13) | **<0.001** | 0.06 (-0.03 – 0.14) | 0.207 | -0.03 (-0.10 – 0.04) | 0.375 | -0.15 (-0.23 – -0.07) | **<0.001** | -0.02 (-0.03 – -0.01) | **0.002** |
| Sex (male) | -0.01 (-0.14 – 0.11) | 0.821 | -0.60 (-0.67 – -0.54) | **<0.001** | 0.05 (-0.00 – 0.10) | 0.073 | -0.07 (-0.13 – -0.00) | **0.036** | -0.00 (-0.01 – 0.01) | 0.883 |
| Age (in years) | 0.01 (0.01 – 0.02) | **<0.001** | 0.01 (0.01 – 0.01) | **<0.001** | 0.00 (0.00 – 0.01) | **<0.001** | -0.01 (-0.02 – -0.01) | **<0.001** | -0.00 (-0.00 – -0.00) | **<0.001** |
| Season (wet) | 0.16 (0.02 – 0.31) | **0.029** | 0.18 (0.13 – 0.23) | **<0.001** | -0.09 (-0.13 – -0.04) | **<0.001** | -0.15 (-0.19 – -0.10) | **<0.001** | -0.01 (-0.02 – -0.00) | **0.002** |
| High WBC | 0.42 (0.16 – 0.69) | **0.002** | 0.23 (0.14 – 0.32) | **<0.001** | 1.82 (1.74 – 1.90) | **<0.001** | 0.99 (0.90 – 1.07) | **<0.001** | 0.05 (0.04 – 0.07) | **<0.001** |
| **Random Effects** | | | | | | | | | | |
| Residual σ^2^ | 0.87 | | 0.64 | | 0.63 | | 0.66 | | 0.01 | |
| τ_00_ | 0.09 _pid_ | | 0.22 _pid_ | | 0.09 _pid_ | | 0.15 _pid_ | | 0.00 _pid_ | |
|  | 0.03 _community_id_ | | 0.03 _community_id_ | | 0.03 _community_id_ | | 0.06 _community_id_ | | 0.00 _community_id_ | |
| ICC | 0.13 | | 0.28 | | 0.16 | | 0.25 | | 0.26 | |
| N | 894 _pid_ | | 1255 _pid_ | | 1261 _pid_ | | 1261 _pid_ | | 1261 _pid_ | |
|  | 68 _community_id_ | | 80 _community_id_ | | 80 _community_id_ | | 80 _community_id_ | | 80 _community_id_ | |
| Observations | 1011 | | 5921 | | 6078 | | 6063 | | 6054 | |
| Marginal R^2^ / Conditional R^2^ | 0.053 / 0.172 | | 0.112 / 0.363 | | 0.235 / 0.355 | | 0.111 / 0.330 | | 0.027 / 0.277 | |

Supplemental Table 2a. Mixed effects linear regressions assessing the association between *APOE* genotype and measures of innate immune function. The main predictor of interest, *APOE4*, is a binary variable that represents having at least a single copy of the *E4* allele. All dependent variables were transformed and centered for analyses. Random effects are included for individual and community residence. Results are reported as standardized betas (2.5% - 97.5% confidence intervals)

|  | **Leukocytes** | | **Lymphocytes** | | **Hemoglobin** | |
| --- | --- | --- | --- | --- | --- | --- |
| *Predictors* | *β ( 95%CI )* | *p* | *β ( 95%CI )* | *p* | *β ( 95%CI )* | *p* |
| (Intercept) | 0.30 (0.16 – 0.43) | **<0.001** | 0.64 (0.48 – 0.80) | **<0.001** | 0.75 (0.58 – 0.92) | **<0.001** |
| *APOE4* | -0.09 (-0.15 – -0.02) | **0.012** | 0.01 (-0.07 – 0.09) | 0.878 | -0.04 (-0.13 – 0.06) | 0.442 |
| Sex (male) | -0.04 (-0.09 – 0.02) | 0.182 | -0.13 (-0.19 – -0.06) | **<0.001** | 0.71 (0.63 – 0.78) | **<0.001** |
| Age (in years) | -0.01 (-0.01 – -0.00) | **<0.001** | -0.01 (-0.01 – -0.01) | **<0.001** | -0.02 (-0.02 – -0.01) | **<0.001** |
| Season (wet) | -0.17 (-0.20 – -0.13) | **<0.001** | -0.15 (-0.20 – -0.10) | **<0.001** | -0.28 (-0.32 – -0.24) | **<0.001** |
| High WBC | 2.01 (1.94 – 2.08) | **<0.001** | 1.02 (0.93 – 1.11) | **<0.001** | -0.14 (-0.22 – -0.07) | **<0.001** |
| **Random Effects** | | | | | | |
| Residual σ^2^ | 0.48 | | 0.70 | | 0.52 |  |
| τ_00_ | 0.12 _pid_ | | 0.17 _pid_ | | 0.32 _pid_ |  |
|  | 0.04 _community_id_ | | 0.02 _community_id_ | | 0.02 _community_id_ |  |
| ICC | 0.24 | | 0.21 | | 0.39 |  |
| N | 1265 _pid_ | | 1261 _pid_ | | 1264 _pid_ |  |
|  | 80 _community_id_ | | 80 _community_id_ | | 80 _community_id_ |  |
| Observations | 6331 | | 6069 | | 6301 |  |
| Marginal R^2^ / Conditional R^2^ | 0.319 / 0.485 | | 0.099 / 0.286 | | 0.176 / 0.501 |  |

Supplemental Table 2b. Mixed effects linear regressions assessing the association between *APOE* genotype and other measures of immune function. The main predictor of interest, *APOE4*, is a binary variable that represents having at least a single copy of the *E4* allele. All dependent variables were transformed and centered for analyses. Random effects are included for individual and community residence. Results are reported as standardized betas (2.5% - 97.5% confidence intervals).

|  | **BMI** | | | **Total Cholesterol** | | | **LDL** | | | **HDL** | | | **Ox-LDL** | | | **Triglycerides** | | |
| --- | --- | --- | --- | --- | --- | --- | --- | --- | --- | --- | --- | --- | --- | --- | --- | --- | --- | --- |
| *Predictors* | | *β ( 95%CI )* | *p* | | *β ( 95%CI )* | *p* | | *β ( 95%CI )* | *p* | | *β ( 95%CI )* | *p* | | *β ( 95%CI )* | *p* | | *β ( 95%CI )* | *p* |
| (Intercept) | | 0.37 (0.20 – 0.54) | **<0.001** | | -0.46 (-0.70 – -0.22) | **<0.001** | | -0.39 (-0.63 – -0.15) | **0.002** | | -0.60 (-0.84 – -0.36) | **<0.001** | | 0.40 (0.03 – 0.78) | **0.034** | | -0.06 (-0.31 – 0.19) | 0.633 |
| *APOE4* | | 0.15 (0.02 – 0.28) | **0.027** | | 0.16 (0.04 – 0.27) | **0.008** | | 0.08 (-0.04 – 0.19) | 0.193 | | 0.05 (-0.06 – 0.17) | 0.348 | | 0.17 (0.01 – 0.33) | **0.043** | | 0.07 (-0.05 – 0.19) | 0.272 |
| Sex (male) | | -0.05 (-0.16 – 0.05) | 0.323 | | -0.14 (-0.23 – -0.05) | **0.002** | | -0.19 (-0.28 – -0.10) | **<0.001** | | 0.08 (-0.01 – 0.16) | 0.086 | | -0.09 (-0.22 – 0.04) | 0.157 | | -0.12 (-0.22 – -0.02) | **0.015** |
| Age (years) | | -0.01 (-0.01 – -0.00) | **<0.001** | | 0.01 (0.01 – 0.01) | **<0.001** | | 0.01 (0.01 – 0.01) | **<0.001** | | 0.01 (0.00 – 0.01) | **<0.001** | | -0.00 (-0.01 – 0.00) | 0.156 | | 0.00 (-0.00 – 0.01) | 0.603 |
| Season (wet) | | -0.08 (-0.11 – -0.06) | **<0.001** | | -0.12 (-0.20 – -0.04) | **0.003** | | -0.30 (-0.38 – -0.21) | **<0.001** | | 0.08 (-0.01 – 0.17) | 0.071 | | -0.31 (-0.47 – -0.15) | **<0.001** | | 0.09 (0.01 – 0.16) | **0.021** |
| High WBC | | -0.02 (-0.07 – 0.03) | 0.398 | | -0.02 (-0.17 – 0.14) | 0.819 | | -0.08 (-0.24 – 0.08) | 0.321 | | -0.08 (-0.24 – 0.08) | 0.321 | | 0.05 (-0.20 – 0.30) | 0.698 | | 0.06 (-0.08 – 0.20) | 0.426 |
| **Random Effects** | | | | | | | | | | | | | | | | | | |
| Residual σ^2^ | | 0.19 | | | 0.70 | | | 0.71 | | | 0.83 | | | 0.30 | | | 0.56 | |
| τ_00_ | | 0.84 _pid_ | | | 0.24 _pid_ | | | 0.18 _pid_ | | | 0.14 _pid_ | | | 0.65 _pid_ | | | 0.39 _pid_ | |
|  | | 0.03 _community_id_ | | | 0.09 _community_id_ | | | 0.11 _community_id_ | | | 0.10 _community_id_ | | | 0.11 _community_id_ | | | 0.05 _community_id_ | |
| ICC | | 0.82 | | | 0.32 | | | 0.29 | | | 0.22 | | | 0.71 | | | 0.44 | |
| N | | 1263 _pid_ | | | 1161 _pid_ | | | 1148 _pid_ | | | 1160 _pid_ | | | 894 _pid_ | | | 1167 _pid_ | |
|  | | 80 _community_id_ | | | 71 _community_id_ | | | 71 _community_id_ | | | 71 _community_id_ | | | 68 _community_id_ | | | 71 _community_id_ | |
| Observations | | 6327 | | | 2538 | | | 2352 | | | 2438 | | | 1011 | | | 2589 | |
| Marginal R^2^ / Conditional R^2^ | | 0.009 / 0.819 | | | 0.021 / 0.333 | | | 0.041 / 0.319 | | | 0.012 / 0.232 | | | 0.029 / 0.722 | | | 0.006 / 0.447 | |

Supplemental Table 3. Mixed effects linear regressions assessing the association between *APOE* genotype and measures of lipids and BMI. The main predictor of interest, *APOE4*, is a binary variable that represents having at least a single copy of the *E4* allele. All dependent variables were transformed and centered for analyses. Random effects are included for individual and community residence. Results are reported as standardized betas (2.5% - 97.5% confidence intervals).

|  | Age (in yrs) | | | | | E4 | | | | | Age * E4 | | | | |
| --- | --- | --- | --- | --- | --- | --- | --- | --- | --- | --- | --- | --- | --- | --- | --- |
| *Models* | *β* | *CI* | | | *p* | *β* | | *CI* | | *p* | *β* | | *CI* | | *p* |
| CRP | 0.02 | 0.01 – 0.02 | | | **<0.001** | -0.62 | | -1.44 – 0.21 | | 0.142 | 0.01 | | -0.01 – 0.02 | | 0.454 |
| Sed rate | 0.01 | 0.00 – 0.01 | | | **<0.001** | -0.03 | | -0.40 – 0.34 | | 0.866 | 0.00 | | -0.01 – 0.01 | | 0.607 |
| Eosin : Lymph | -0.01 | -0.01 – -0.01 | | | **<0.001** | -0.44 | | -0.81 – -0.07 | | **0.02** | 0.01 | | -0.00 – 0.01 | | 0.091 |
| Lymphocytes | -0.01 | | -0.01 – -0.01 | **<0.001** | | 0.14 | -0.21 – 0.50 | | 0.425 | | 0.00 | -0.01 – 0.00 | | 0.379 | |
| Eosinophils | -0.02 | -0.02 – -0.01 | | | **<0.001** | -0.40 | | -0.75 – -0.06 | | **0.022** | 0.00 | | -0.00 – 0.01 | | 0.137 |
| Neutrophils | 0.00 | 0.00 – 0.01 | | | **0.002** | -0.25 | | -0.56 – 0.06 | | 0.117 | 0.00 | | -0.00 – 0.01 | | 0.165 |

Supplemental Table 4. Models showing immune function and age associations, including interactive effects with *APOE* status. Results are from mixed effects linear regressions (for full table see Supplemental Table XY), adjusting for sex, season, and a dummy variable used as a proxy for current illness (leukocytes > 12 mm3). Results are reported as standardized betas; *CI* is the 95% confidence interval. All dependent variables were transformed and centered prior to analyses. *APOE* genotype is coded as a categorical variable, binned as individuals that are homozygous *E3* (E3) versus those that have at least one copy of the *E4* allele (E4).

|  | **CRP** | | **Sed Rate** | | **Neutrophils** | |
| --- | --- | --- | --- | --- | --- | --- |
| *Predictors* | *β ( 95% CI )* | *p* | *β ( 95% CI )* | *p* | *β ( 95% CI )* | *p* |
| (Intercept) | -0.69 (-1.08 – -0.30) | **0.001** | -0.81 (-1.04 – -0.58) | **<0.001** | -0.38 (-0.58 – -0.18) | **<0.001** |
| Total Cholesterol | -0.14 (-0.21 – -0.06) | **<0.001** | -0.06 (-0.10 – -0.02) | **0.002** | 0.02 (-0.01 – 0.06) | 0.258 |
| BMI | 0.11 (0.04 – 0.17) | **0.002** | -0.03 (-0.07 – 0.01) | 0.094 | 0.00 (-0.03 – 0.04) | 0.815 |
| Sex (male) | -0.08 (-0.21 – 0.05) | 0.228 | -0.59 (-0.67 – -0.50) | **<0.001** | 0.06 (-0.02 – 0.13) | 0.126 |
| Age (years) | 0.01 (0.01 – 0.02) | **<0.001** | 0.02 (0.01 – 0.02) | **<0.001** | 0.01 (0.00 – 0.01) | **<0.001** |
| Season (wet) | 0.14 (-0.02 – 0.31) | 0.082 | 0.17 (0.09 – 0.25) | **<0.001** | -0.10 (-0.18 – -0.03) | **0.006** |
| High WBC | 0.65 (0.36 – 0.93) | **<0.001** | 0.18 (0.04 – 0.33) | **0.015** | 1.85 (1.71 – 1.98) | **<0.001** |
| Total Cholesterol * BMI | 0.13 (0.07 – 0.18) | **<0.001** | 0.04 (0.01 – 0.08) | **0.008** | 0.02 (-0.01 – 0.05) | 0.155 |
| **Random Effects** | | | | | | |
| Residual σ^2^ | 0.89 | | 0.62 | | 0.60 | |
| τ_00_ | 0.04 _pid_ | | 0.20 _pid_ | | 0.10 _pid_ | |
|  | 0.04 _community_id_ | | 0.05 _community_id_ | | 0.04 _community_id_ | |
| ICC | 0.08 | | 0.29 | | 0.19 | |
| N | 771 _pid_ | | 1144 _pid_ | | 1157 _pid_ | |
|  | 65 _community_id_ | | 71 _community_id_ | | 71 _community_id_ | |
| Observations | 878 | | 2488 | | 2520 | |
| Marginal R^2^ / Conditional R^2^ | 0.084 / 0.160 | | 0.134 / 0.388 | | 0.221 / 0.370 | |

Supplemental Table 5. Mixed effects linear regression models that test direct and interactive effects of total cholesterol and BMI on inflammatory immune markers. Random effects are included for individual and community residence. Dependent and predictor variables of interest (total cholesterol, BMI, and immune markers) were transformed and centered for analyses. Results are reported as standardized betas (2.5% - 97.5% confidence intervals).

|  | **CRP** | | **Sed Rate** | | **Neutrophils** | |
| --- | --- | --- | --- | --- | --- | --- |
| *Predictors* | *β ( 95% CI )* | *p* | *β ( 95% CI )* | *p* | *β ( 95% CI )* | *p* |
| (Intercept) | -0.68 (-1.07 – -0.29) | **0.001** | -0.81 (-1.04 – -0.57) | **<0.001** | -0.38 (-0.59 – -0.18) | **<0.001** |
| LDL Cholesterol | -0.11 (-0.19 – -0.04) | **0.003** | -0.03 (-0.07 – 0.01) | 0.172 | 0.03 (-0.01 – 0.07) | 0.113 |
| BMI | 0.08 (0.00 – 0.15) | **0.039** | -0.03 (-0.07 – 0.01) | 0.183 | 0.01 (-0.03 – 0.04) | 0.763 |
| Sex (male) | -0.07 (-0.20 – 0.06) | 0.295 | -0.58 (-0.67 – -0.50) | **<0.001** | 0.06 (-0.02 – 0.13) | 0.152 |
| Age (years) | 0.01 (0.01 – 0.02) | **<0.001** | 0.02 (0.01 – 0.02) | **<0.001** | 0.01 (0.00 – 0.01) | **<0.001** |
| Season (wet) | 0.10 (-0.06 – 0.27) | 0.218 | 0.17 (0.08 – 0.25) | **<0.001** | -0.05 (-0.13 – 0.02) | 0.167 |
| High WBC | 0.65 (0.36 – 0.94) | **<0.001** | 0.16 (0.01 – 0.31) | **0.036** | 1.83 (1.69 – 1.97) | **<0.001** |
| LDL Cholesterol * BMI | 0.15 (0.09 – 0.21) | **<0.001** | 0.03 (0.00 – 0.06) | 0.087 | 0.01 (-0.03 – 0.04) | 0.713 |
| **Random Effects** | | | | | | |
| Residual σ^2^ | 0.89 | | 0.63 | | 0.60 | |
| τ_00_ | 0.05 _pid_ | | 0.21 _pid_ | | 0.11 _pid_ | |
|  | 0.04 _community_id_ | | 0.05 _community_id_ | | 0.04 _community_id_ | |
| ICC | 0.09 | | 0.29 | | 0.20 | |
| N | 765 _pid_ | | 1130 _pid_ | | 1144 _pid_ | |
|  | 65 _community_id_ | | 71 _community_id_ | | 71 _community_id_ | |
| Observations | 873 | | 2305 | | 2344 | |
| Marginal R^2^ / Conditional R^2^ | 0.083 / 0.163 | | 0.128 / 0.385 | | 0.219 / 0.373 | |

Supplemental Table 6. Mixed effects linear regression models that test direct and interactive effects of LDL cholesterol and BMI on inflammatory immune markers. Random effects are included for individual and community residence. Dependent and predictor variables of interest (LDL cholesterol, BMI, and immune markers) were transformed and centered for analyses. Results are reported as standardized betas (2.5% - 97.5% confidence intervals).

|  | **CRP** | | **Sed Rate** | | **Neutrophils** | |
| --- | --- | --- | --- | --- | --- | --- |
| *Predictors* | *β ( 95% CI )* | *p* | *β ( 95% CI )* | *p* | *β ( 95% CI )* | *p* |
| (Intercept) | -0.98 (-1.33 – -0.64) | **<0.001** | -0.66 (-1.00 – -0.32) | **<0.001** | -0.53 (-0.81 – -0.24) | **<0.001** |
| Oxidized LDL | -0.00 (-0.07 – 0.06) | 0.886 | 0.00 (-0.06 – 0.06) | 0.908 | -0.01 (-0.06 – 0.04) | 0.710 |
| BMI | 0.10 (0.03 – 0.16) | **0.002** | 0.02 (-0.04 – 0.08) | 0.541 | 0.01 (-0.04 – 0.06) | 0.829 |
| Sex (male) | -0.01 (-0.13 – 0.12) | 0.920 | -0.48 (-0.59 – -0.37) | **<0.001** | 0.09 (-0.00 – 0.19) | 0.063 |
| Age (years) | 0.02 (0.01 – 0.02) | **<0.001** | 0.01 (0.01 – 0.02) | **<0.001** | 0.01 (0.01 – 0.02) | **<0.001** |
| Season (wet) | 0.16 (0.02 – 0.31) | **0.026** | -0.18 (-0.33 – -0.02) | **0.026** | 0.18 (0.05 – 0.31) | **0.007** |
| High WBC | 0.51 (0.25 – 0.78) | **<0.001** | 0.25 (0.01 – 0.49) | **0.043** | 1.92 (1.71 – 2.13) | **<0.001** |
| Oxidized LDL * BMI | 0.14 (0.08 – 0.19) | **<0.001** | -0.04 (-0.09 – 0.01) | 0.137 | 0.02 (-0.02 – 0.06) | 0.352 |
| **Random Effects** | | | | | | |
| Residual σ^2^ | 0.87 | | 0.56 | | 0.54 | |
| τ_00_ | 0.06 _pid_ | | 0.15 _pid_ | | 0.04 _pid_ | |
|  | 0.03 _community_id_ | | 0.13 _community_id_ | | 0.07 _community_id_ | |
| ICC | 0.10 | | 0.33 | | 0.16 | |
| N | 893 _pid_ | | 831 _pid_ | | 890 _pid_ | |
|  | 68 _community_id_ | | 68 _community_id_ | | 68 _community_id_ | |
| Observations | 1010 | | 945 | | 1005 | |
| Marginal R^2^ / Conditional R^2^ | 0.078 / 0.169 | | 0.093 / 0.396 | | 0.274 / 0.393 | |

Supplemental Table 7. Mixed effects linear regression models that test direct and interactive effects of oxidized LDL and BMI on inflammatory immune markers. Random effects are included for individual and community residence. Dependent and predictor variables of interest (oxidized LDL, BMI, and immune markers) were transformed and centered for analyses. Results are reported as standardized betas (2.5% - 97.5% confidence intervals).

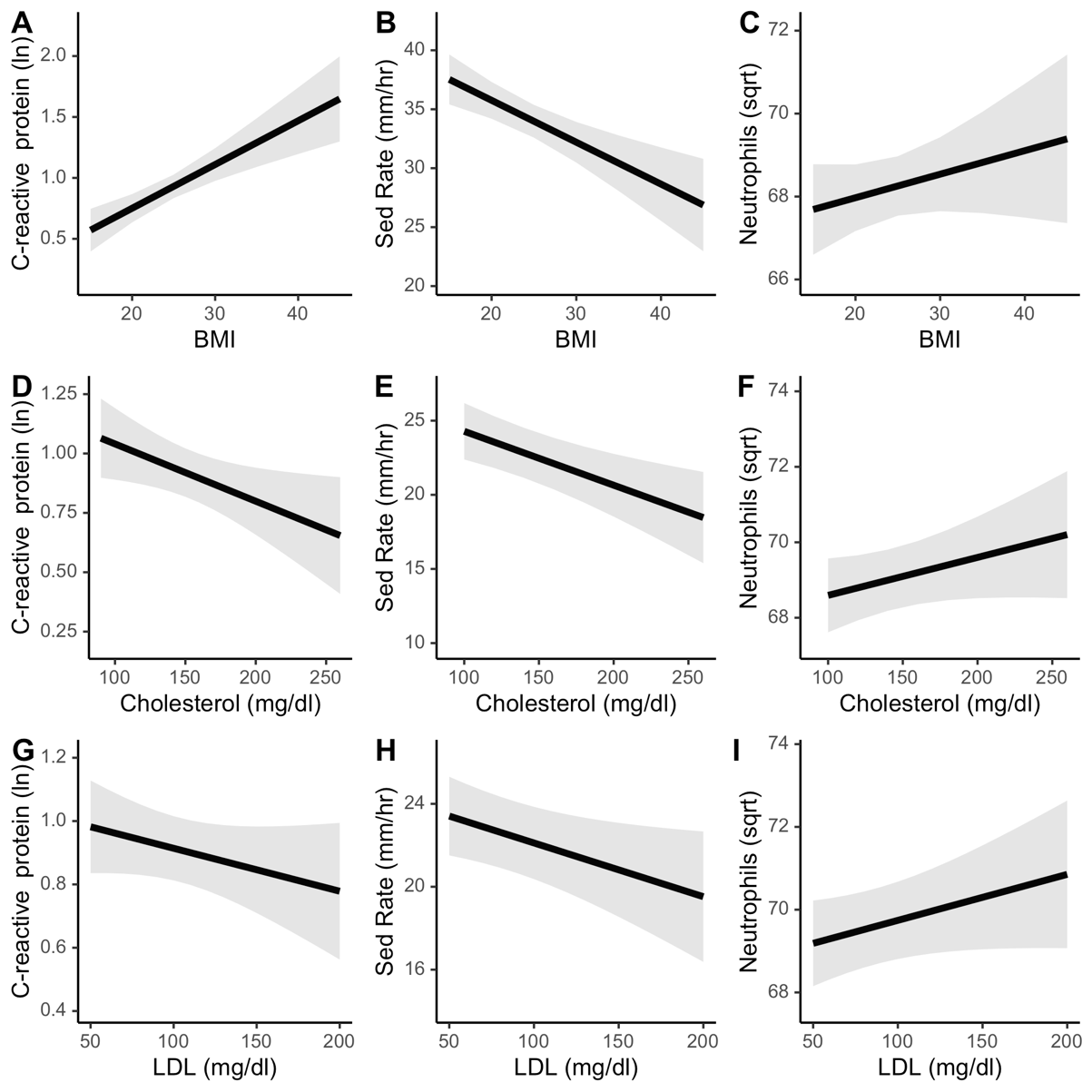

Supplemental Figure 1. This figure shows independent associations between BMI and cholesterols with markers of inflammation.
